# Dual-color volumetric imaging of neural activity of cortical columns

**DOI:** 10.1101/504233

**Authors:** Shuting Han, Weijian Yang, Rafael Yuste

**Affiliations:** Neurotechnology Center, Department of Biological Sciences, Columbia University, New York, NY 10027, USA; Current address: Department of Electrical and Computer Engineering, University of California, Davis, CA 95616, USA.

**Keywords:** volumetric imaging, two-photon calcium imaging, beam multiplexing, wavelength multiplexing, wavefront shaping, spatial light modulator, ensembles

## Abstract

To capture the emergent properties of neural circuits, high-speed volumetric imaging of neural activity at cellular resolution is desirable. But while conventional two-photon calcium imaging is a powerful tool to study population activity *in vivo*, it is restrained to two-dimensional planes. Expanding it to 3D while maintaining high spatiotemporal resolution appears necessary. Here, we developed a two-photon microscope with dual-color laser excitation that can image neural activity in a 3D volume. We imaged the neuronal activity of primary visual cortex from awake mice, spanning from L2 to L5 with 10 planes, at a rate of 10 vol/sec, and demonstrated volumetric imaging of L1 long-range PFC projections and L2/3 somatas. Using this method, we map visually-evoked neuronal ensembles in 3D, finding a lack of columnar structure in orientation responses and revealing functional correlations between cortical layers which differ from trial to trial and are missed in sequential imaging. We also reveal functional interactions between presynaptic L1 axons and postsynaptic L2/3 neurons. Volumetric two-photon imaging appears an ideal method for functional connectomics of neural circuits.

## Highlights

- Dual-color two-photon volumetric imaging of record numbers of neurons across cortical columns *in vivo*
- Neuronal ensembles span across layers and are not detected in sequential imaging
- Lack of columnar structures in orientation response in V1
- Functional interaction between axonal projections and local population

## Introduction

High-speed volumetric imaging of neural activity at cellular resolution is an important method to decipher the function of microcircuits at a population level. As the mammalian cortex is organized into layers, the coordinated activity of neurons within and across layers likely contributes to emergent functional properties of circuit, making it necessary to measure neuronal activity in three dimensions in order to capture them (Alivisatos et al., 2012). Calcium imaging provides a powerful tool for recording the activity from a large population of neuron *in vivo* (Tian et al., 2009; Yuste and Katz, 1991). In combination with two-photon imaging, it allows the observation of population activity from deep tissues (Denk et al., 1990; Helmchen and Denk, 2005; Yuste and Denk, 1995). However, conventional two-photon calcium imaging is constrained to imaging a single two-dimensional plane. To extend it to a three-dimensional volume while still maintaining cellular resolution and high temporal resolution, multiple strategies have been developed (Yang and Yuste, 2017). Fast z-scan devices such as spatial light modulators (SLMs) and electric tunable lenses (ETLs) are capable of switching focus at high speed over a relatively large depth range (up to ∼500 μm), and have been demonstrated for fast sequential volumetric imaging (Grewe et al., 2011; Yang et al., 2016). In addition, as holographic devices, SLMs can be used to generate multiple beamlets to simultaneously image multiple planes across >500 µm, with their signals demixed by statistical algorithms (Pnevmatikakis et al., 2016; Yang et al., 2016). Other volumetric imaging approaches such as remote focusing introduces an auxiliary objective and a scan mirror that allows fast switching of focal planes (Botcherby et al., 2008, 2012).

Here we developed a new hybrid approach combining excitation-wavelength multiplexing and fast z-scan devices. We labeled superficial and deep neuronal population with two fluorophores of different colors, and simultaneously excited each population with a different laser. To minimize scattering effect, we chose GCaMP6 (Chen et al., 2013) for superficial layers (layer 2/3), and the red-shifted jRGECO1b (Dana et al., 2016) for deep layers (layer 5) (Dana et al., 2016; Tian et al., 2009). We used an ETL for fast sequential z-scanning in superficial layers and an SLM for deep layers through wavefront shaping simultaneously. With these, we demonstrated imaging 10 planes over 450 μm that spans from layer 1 to layer 5 in primary visual cortex (V1) of awake mice, while maintaining a high temporal resolution of 10 vol/sec. We further demonstrated volumetric imaging of layer 1 local dendrites with layer 2/3 somas, and long-range projections from PFC in layer 1 of V1 with local layer 2/3 population. We identified visually-evoked ensembles in 3D, showed a lack of columnar structures in the responses, and revealed that correlation between cortical layers varies from trial to trial which is missed by sequential imaging.

## Results

### Dual-color two-photon microscope

Our microscope consists of two beam paths with two separate two-photon lasers, exciting green and red calcium indicators, correspondingly. The beam path for green indicator (GCaMP6) includes an ultrafast laser (920 nm), a telescope that expands the beam to fill the ETL, and an ETL for fast sequential defocusing. The beam path for red indicator (jRGECO), based on a previous design (Yang et al., 2016), has an ultrafast holographic focal planes. The default focal plane of the SLM path is separated from the ETL path by 200 μm deeper, implemented by an offset lens at the conjugate plane of the imaging plane (Figure 1A). Placing the 1064 nm laser with red indicator in the deep layers benefits from less scattering of longer wavelength excitation and emission. The two lasers combine at a dichroic mirror, and are then scanned by a resonant scanner and a galvanometric scanner, and simultaneously excite the sample at different depths. The emitted fluorescence is separated by another dichroic mirror, and collected by two PMTs with filters optimized for corresponding fluorophores. Additionally, in order to optimize for large angle scanning, we adopted the lens complex design between the two scanners based on (Stirman et al., 2016) (Figure 1B). For volumetric imaging *in vivo,* two planes of ∼200 μm apart are excited and recorded simultaneously; the dual focal planes are directed to a set of depths that covers two separate volumes in synchronous through ETL and SLM (Figure 1C).

**Figure 1.**
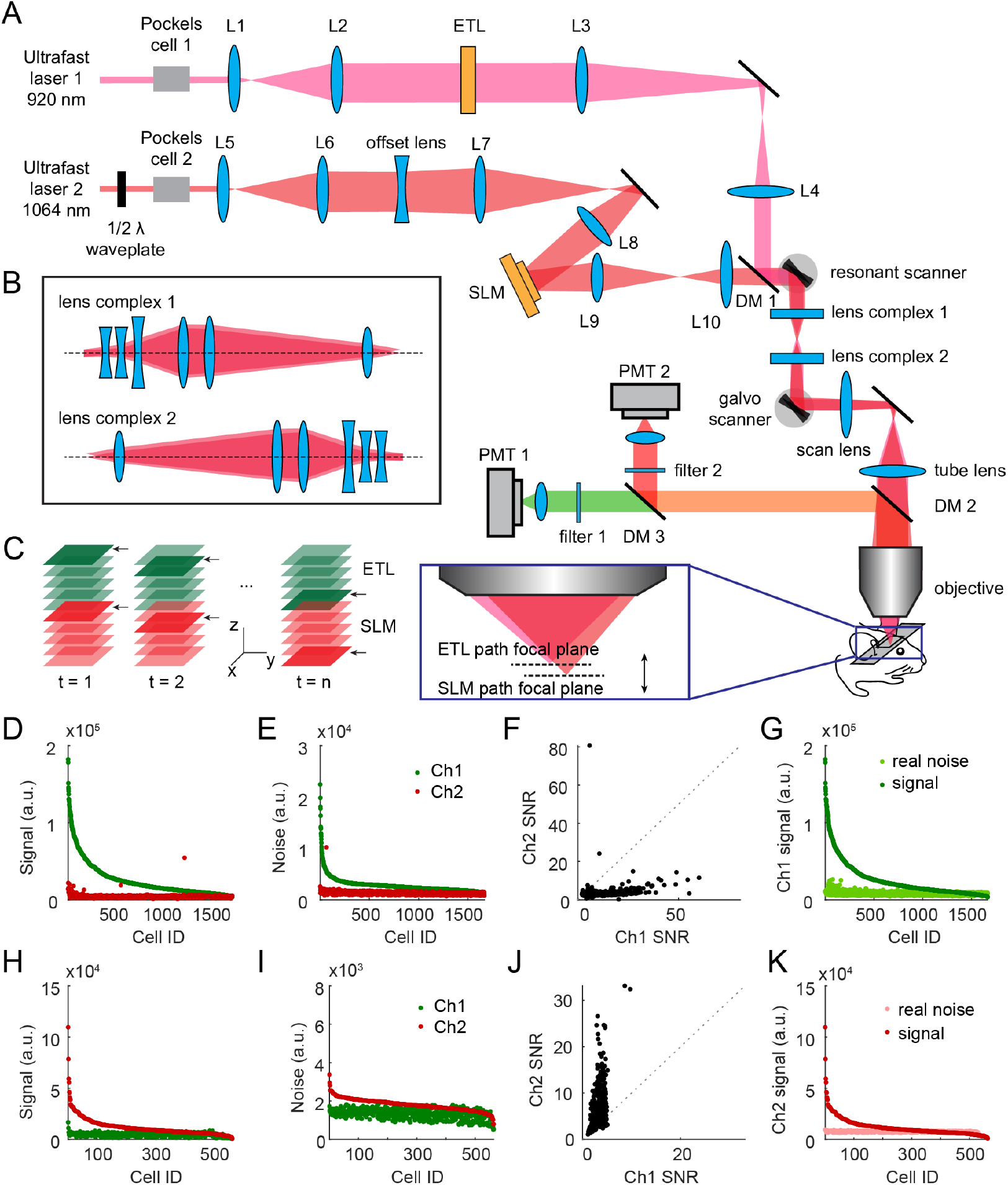
Dual-color two-photon volumetric imaging microscope. (A) Microscope design diagram. Two lasers at 920 nm and 1064 nm are expanded and modulated by an ETL unit and an SLM unit, correspondingly, then combined through a dichroic mirror and pass through a resonant scanner and a galvo scanner before exciting the sample through objective lens. Emission fluorescence is collected through two separate PMTs. The SLM path is also equipped with an offset lens that separates its focal plane (200 µm deeper) from that of the ETL path. (B) Details of the lens complex design. (C) Diagram of volumetric imaging. Two planes are excited and recorded at the same time: the shallower one from the ETL path, and the deeper one from the SLM path. The imaging depth of the dual planes cycles over time to record a 3D volume. (D-F) Measured signal (D), noise (E), and signal-to-noise ratio (SNR, F) from both green (Ch1) and red (Ch2) PMTs, with only 920 nm laser on. (G) Quantified signal strength with real noise in green channel (Ch1), with only 920 nm laser on. Real noise is computed as noise in (E) (green dots) + signal in (H) (red dots) + noise in (I) (red dots). (H-J) Measured signal (H), noise (I), and SNR (J) from both green (Ch1) and red (Ch2) PMTs, with only 1064 nm laser on. (K) Quantified signal strength with real noise in red channel (Ch2), with only 1064 nm laser on. Real noise is computed as noise in (H) (red dots) + signal in (D) (green dots) + noise in (E) (green dots). (n = 3 experiments)

To optimize for deep layer imaging and to compensate for the optical aberrations in the system, we also implemented adaptive optics (AO) with the SLM in the 1064 nm excitation path. We modeled the wavefront aberrations with a combination of Zernike polynomials aberration modes, measured their coefficients using fluorescent beads (Love, 1997), and then corrected for the wavefront using the SLM (Figure S1). AO improves both the target intensity and the PSF over ±200 μm defocus range, reaching a minimum FWHM of 6 μm (Figure S2B, S2D). Wavefront correction for the ETL path is less critical as it images superficial layers. It has a minimum FWHM of 8 μm (Figure S2A, S2C).

To ensure that the signals we recorded from the simultaneous dual planes do not interfere, we characterized the cross-channel contamination of our system. We imaged mice V1 *in vivo* by turning on the 920 nm laser only, or the 1064 nm laser only, while recording signals from both PMTs simultaneously. In this case, signals from the non-exciting channel represent potential contamination. We then analyzed the signal and noise from both channels with single laser excitation. For both single laser excitations, the desired signal (green for 920 nm excitation in Figure 1D, red for 1064 nm excitation in Figure 1H) are much higher than the cross-channel contamination (red in Figure 1D, green in Figure 1H), while the noise exhibits similar patterns (Figure 1E, 1I). Overall, the signal to noise ratio (SNR) is much higher in the desired channel (Figure 1F, 1J). We further estimated the real noise from each channel, assuming that both lasers are exciting, by adding up the noise from desired channel with corresponding laser, and bleached through signal and noise in the same channel produced by the other laser. Under this estimation, the signal is still higher than the estimated real noise in both channels (Figure 1G, 1K). We conclude that our system has minimal cross-channel contamination and is optimized for simultaneous dual-color imaging.

### *In vivo* volumetric imaging of cortical columns

We applied our system to image the cortical activity of neurons from awake mice V1. We labeled the V1 neuron population with the green GCaMP6s (Chen et al., 2013) and the red jRGECO1b (Dana et al., 2016) through viral vectors. We used the ETL beam path to image GCaMP6s with 5 planes spanning from 150 μm to 350 μm in upper layers, and the SLM beam path to image jRGECO1b with 5 planes spanning from 400 μm to 600 μm in lower layers, all spaced with intervals of 50 μm (Figure 2A). This wavelength multiplexing strategy with two beam paths together achieved a total of 10 imaging planes across 450 μm with a standard field of view (FOV) of around 500 μm × 500 μm at each plane, covering from the top of layer 2 through layer 5, at a volume rate of 10.4 vol/sec. In the example shown in Figure 2A, we recorded the spontaneous activity over a 10-minute period from a population of 1497 cells in total.

**Figure 2.**
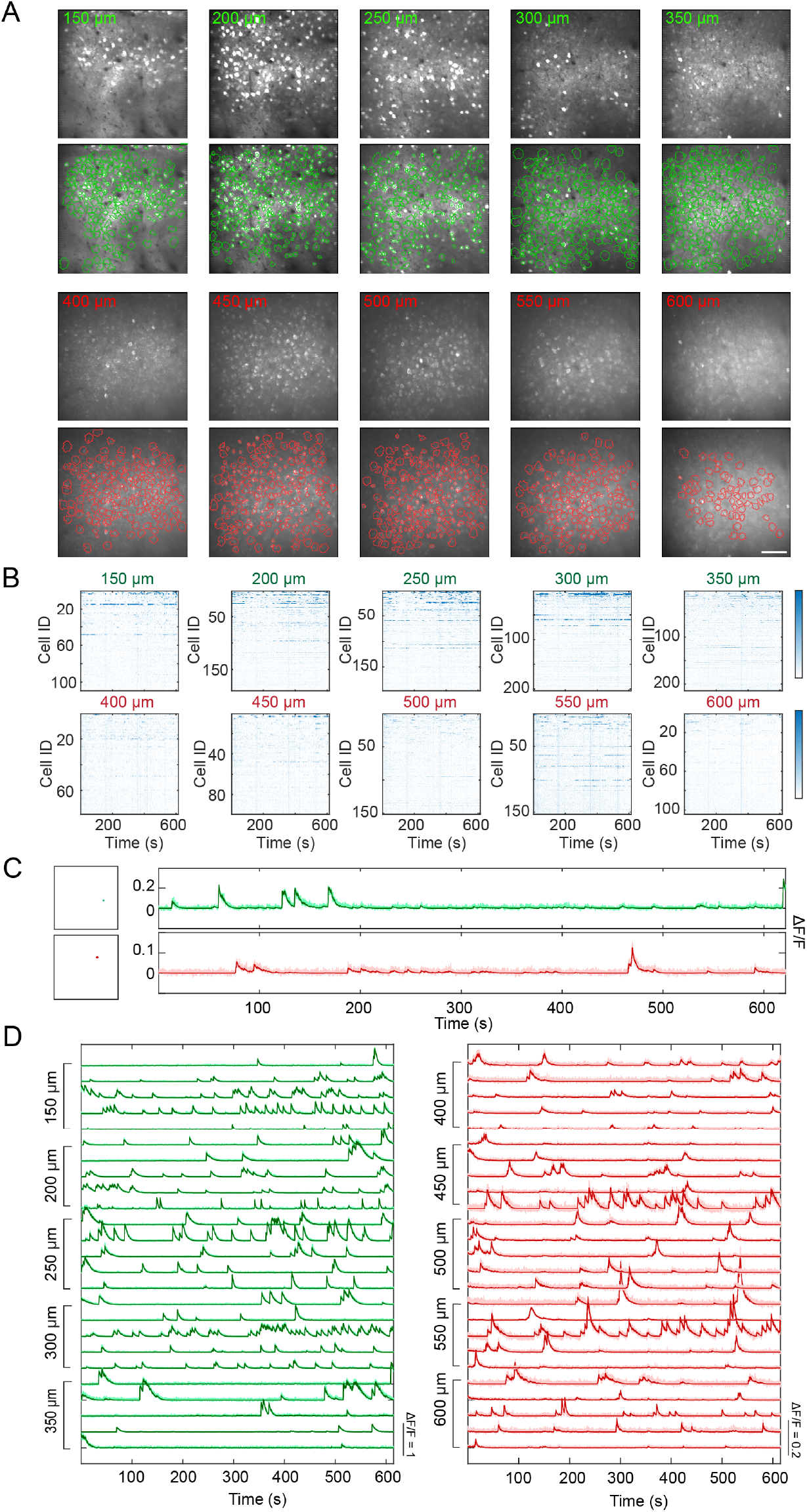
*In vivo* volumetric imaging of cortical columns. (A) Row 1: average images of recorded planes in Ch1 with 920 nm laser, recorded from 150 μm to 350 μm, with a spacing of 50 μm. Row 2: ROI contours extracted by the CNMF algorithm, overlaid with average images. Row 3: average images of recorded planes in Ch2 with 1064 nm laser, recorded from 400 μm to 600 μm, with a spacing of 50 μm. Row 4: ROI contours extracted by the CNMF algorithm, overlaid with average images. Scale bar: 100 μm. (B) Raw ΔF/F traces from all extracted ROIs in each plane, over 10 min spontaneous activity. (C) Two examples of raw (light color) and deconvolved (dark color) traces, from 920 nm and 1064 nm path. (D) Example traces from each plane. Light color: raw traces. Dark color: deconvolved traces.

We then extracted the fluorescence traces from every neuron by a modified version of a constrained nonnegative matrix factorization (CNMF) algorithm (Pnevmatikakis et al., 2016). This version takes manual initialization of neuron locations, and the original CNMF algorithm further automatically optimizes the spatial components (shape of each potential neuron), extracts the raw fluorescence (Figure 2C-D, light traces), filters out the noise, and calculates the deconvolved traces which represents the noiseless estimation of firing probability (Figure 2C-D, dark traces). Extracted traces and neuronal region of interests (ROIs) are manually selected before further analysis. To exclude cross-channel contaminations from particularly “bright” neurons, trace pairs that are highly correlated and come from laterally overlapping ROIs in the simultaneously recorded dual planes (e.g. 150 μm and 400 μm planes) are kept using only the neuron with the highest SNR (Methods). Figure 2A shows examples of extracted (ROIs) from the 10 planes after the above pre-processing steps, and Figure 2B displays their raw traces.

### Orientation selective cells in cortical columns

While conventional two-photon microscopes can image from one cortical layer at a time, our microscope provides a powerful tool for studying neural circuit dynamics across multiple cortical layers with high spatial temporal resolution. To demonstrate this, we recorded visually evoked activity from V1 volumes covering both L2/3 and L5 simultaneously while presenting drifting gratings of 8 directions to the animals. It has been shown that subsets of mouse V1 population are tuned to orientation or direction of drifting gratings (Niell and Stryker, 2008; Rochefort et al., 2011). We indeed identified robust orientation selective cells across the cortical column, in both L2/3 and L/5 (Figure 3A-C). For each orientation, we observed an average of 5%-8% orientation selective cells, in agreement with the characterization previously done by two-photon imaging of L2/3 (Rochefort et al., 2011), while L5 shows less orientation tuned cells of 3%-7%, supporting previous results with extracellular recordings (Niell and Stryker, 2008) (Figure 3D, n = 4 mice, 7 FOVs).

### Visually-evoked neuronal ensembles span superficial and deep layers

In a neuronal circuits, individual neurons cooperate to form larger ensembles of neurons that are functionally correlated. This emergent property of a population, rather than single neurons, is considered to be the functional units during sensory, behavioral and cognitive processes (Carrillo-Reid et al., 2017a; Cossart et al., 2003; Luczak et al., 2007; Mao et al., 2001; Miller et al., 2014; Yuste, 2015). One advantage of our microscope is that we can image multiple cortical layers almost simultaneously, which enables us to define and study cortical ensembles across layers based on the correlation structure of the population. Several computational approaches have been proposed for ensemble detection (Avitan et al., 2017; Carrillo-Reid et al., 2015, 2017b; Lopes-dos-Santos et al., 2013); since we record from a relatively large population of neurons, we chose to use a fast graph-based community detection method [Louvain method (Blondel et al., 2008)] which aims at maximizing modularity measurement. To detect stable ensemble, we combined the Louvain method with consensus clustering that finds the best agreement between repetitions (Lancichinetti and Fortunato, 2012). Here we aimed to find visually-evoked ensembles due to their clear functional correlation with the visual stimulus. We constructed similarity matrices from population activity during visual stimulation, detected and cross-validated neuronal ensembles with the hybrid approach of Louvain method and consensus clustering (Figure 4A; Methods), then evaluated the decoding performance of each ensemble against each visual stimulus. To do this, we generated population vectors from the ensembles, then calculated the cosine similarity between the population vectors and real data, then computed the standard ROC (receiver operating characteristic) curve and AUC (area under curve). We defined visually-evoked ensembles as ensembles that are predictive of visual stimulus with an empirically defined threshold (Figure 4C). For simplicity, we combined pairs of orientations that are shown in opposite directions, resulting ensembles for 4 orientations (Figure 4A-C). The detected ensembles exhibit higher decoding performance than random sampled controls, as well as higher internal pairwise correlation (Figure 4D-E; n = 16 ensembles; AUC control 0.500 ± 0.011 [SEM], ensemble 0.663 ± 0.051 p < 0.001; correlation control 0.092 ± 0.017, ensemble 0.478 ± 0.078, p < 0.001, Wilcoxon signed rank test), indicating their coherent emergent activity as a group.

**Figure 3.**
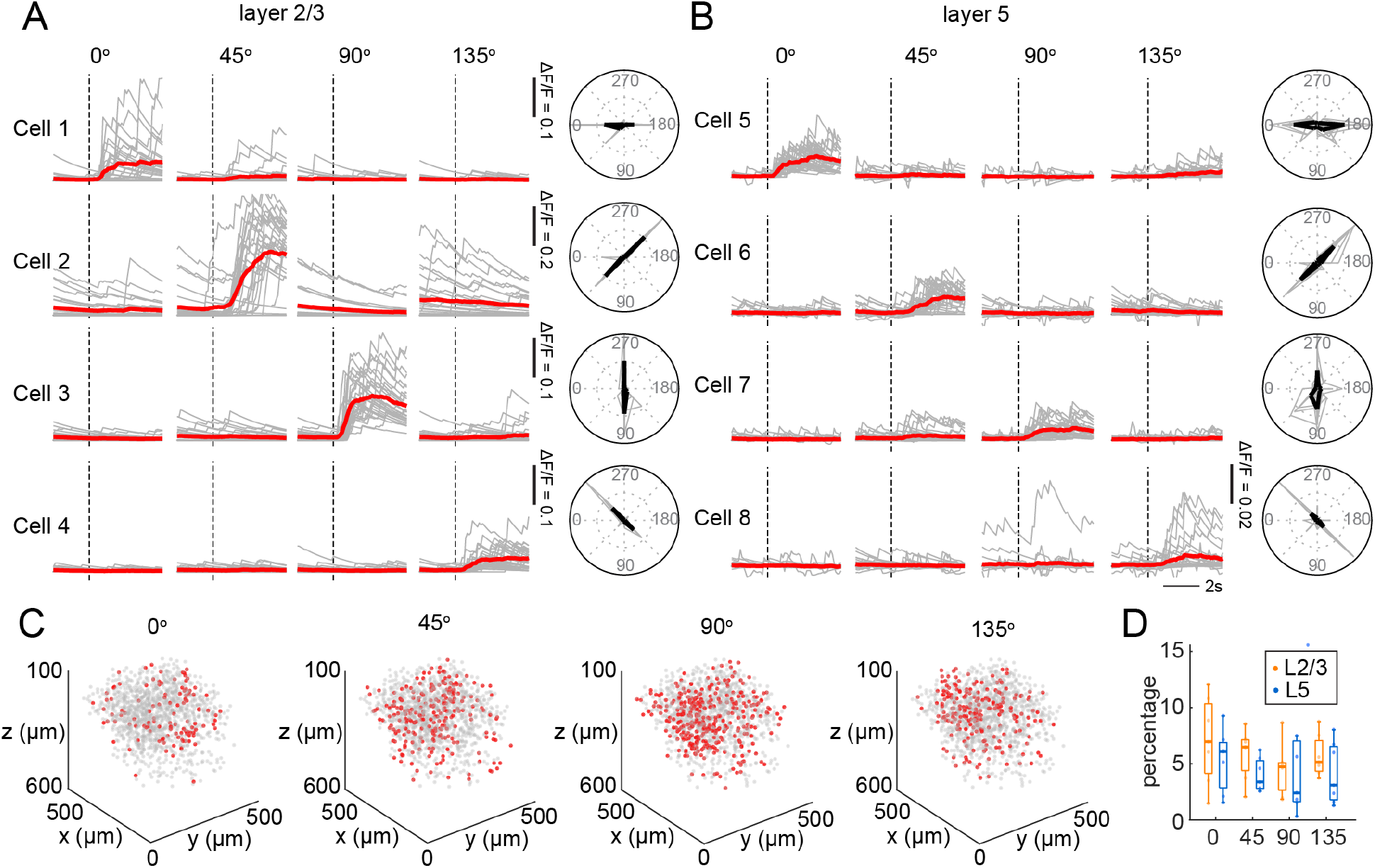
Orientation tuning cells in V1 columns. (A-B) Fluorescence traces (left) and polar graphs (right) of example cell that are selective to 0º, 45º, 90º, and 135º drifting gratings in layer 2/3 (A) and layer 5 (B). (C) 3D distribution of orientation selective cells in the imaged cortical column from an example dataset. (D) Percentage of orientation selective cells in layer 2/3 and layer 5 (n = 7 experiments).

We then investigated the correlation structure of the ensembles between L2/3 and L5 using recorded activity during all visual stimulation trials, or using L2/3 activity during first half of all trials and L5 activity during second half of all trials. The former case represents simultaneous volumetric imaging, while the latter case represents sequential imaging sessions of each layer during repetitive trials, and aligning them with trial onsets. To reduce noise in correlation structures, we investigated functional correlations only within identified visually-evoked ensembles. We separated the ensemble constituent cells into L2/3 and L5 subsets, and computed the pairwise correlation between these two subsets during first and second half trials, or during full trials (Figure 4F). Results combined from 6 datasets show that correlation obtained from full trials are higher than those from half trials, and there is a lack of correlation between full trial correlation and half trial correlation (Figure 4G; R^2^ = 0.08; half trials −0.032 ± 0.046, full trials 0.305 ± 0.080, p < 0.001, Wilcoxon signed rank test). Background activity obtained from non-ROI pixels, however, does not differ drastically (half trials −0.005 ± 0.010, full trials 0.084 ± 0.014, p = 0.031, Wilcoxon signed rank test). This reflects the trial-to-trail variations, possibly due to the animals’ endogenous state (Carandini, 2004; Kiani et al., 2015; Kisley and Gerstein, 1999). When studying functional properties of neural circuits, it thus appears key to simultaneously capture the dynamics of the entire population of interest, since otherwise the trial-to-trial variability will dilute the correlational structure of the activity.

**Figure 4.**
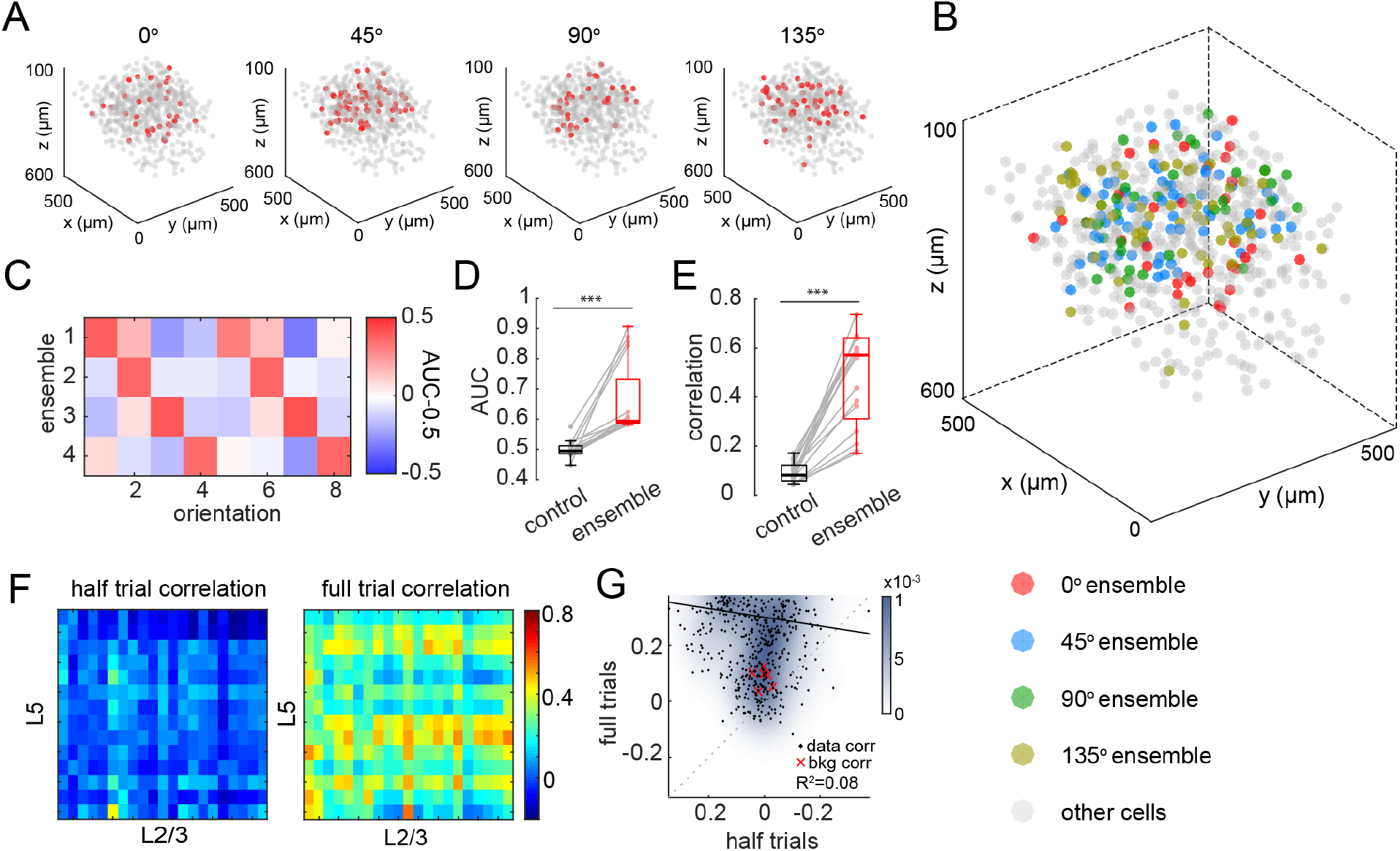
Correlation structure of visually-evoked ensembles. (A) Example of 3D structures of visually-evoked ensembles in the imaged cortical column identified with Louvain method and consensus clustering. (B) 3D view of all visually-evoked ensembles, from the same ensembles as (A). (C) Prediction performance of the example ensembles, of all directions. Color represents AUC - 0.5; red color represents high prediction performance. (D) Statistics of ensemble prediction performance, compared with random controls (p<0.001). Note the y axis represents AUC; 0.5 on AUC axis represents chance level. (E) Statistic of average correlation within ensembles, compared with random controls (p<0.001). Random controls were generated by random sample subsets of the population with the same number of neurons in corresponding ensembles, for 10 times each ensemble. (F) Example correlation structure between ensemble cells in layer 2/3 and layer 5, using the first and second half of visual stimulus trials (left), and using the entire trials (right). (G) Scatter plot of pairwise correlation between layer 2/3 and layer 5 ensemble cells from half trials and full trials. Dashed line represents x = y; black dots represent data point correlations; red crosses represent background correlation from Ch1 and Ch2 in all experiments; black line represents the least-square linear regression result. (n = 7 experiments, 16 ensembles).

### Lack of correlated columnar structures in mouse V1

The visual cortex of some mammalian species is organized into a columnar spatial map where neurons that has similar functional properties such as orientation preference are spatially close to each other (Bonhoeffer and Grinvald, 1991; Hubel and Wiesel, 1962). However, using 2D two-photon calcium imaging, the visual cortex of rodents have been characterized as having a salt-and-pepper structure, where neurons with similar functional properties are intermingled (Ohki and Reid, 2007; Ohki et al., 2005). At the same time, recent studies have reported the existence of narrow (∼40-120 μm diameter) columns neurons with similar tuning properties in rodent V1 (Li et al., 2012; Ringach et al., 2016; Yu et al., 2009). To investigate this controversy, we applied novel our volumetric method, since we could not only analyze the orientation preference map in 3D (Figure 5B), but also extend the analysis to the spatial organization of cells that share emergent properties, which are identified as ensembles (Figure 5A), and functional correlation within narrow columns from the entire population. We analyzed the correlation between lateral distance (distance of xy, ignoring depth) of cell pairs and their evoked activity correlation, among all cell pairs (Figure 5C, left), among visually-evoked ensembles (Figure 5C, middle), and among orientation selective cells (Figure 5C, right). If columnar structure exists, we expect to see higher correlation in cell pairs that are distributed closer laterally. However, all of the three groups show flat distribution, indicating uniform correlation regardless of lateral distance. We further analyzed the correlation values of cell pairs in narrow columns of 30, 50 and 100 μm during visually evoked activities. Compared with random controls where correlation was calculated between cells within the column and a random set of cells outside of the column, none of the column diameters give significant differences (Figure 5D; n = 6 experiments; Wilcoxon rank-sum test on each correlation bin; statistics done with individual experiments). Our results thus indicate a lack of highly correlated column structure in mouse V1, extending to 3D the original 2D salt and pepper description of orientation responses (Ohki and Reid, 2007; Ohki et al., 2005), but in disagreement with the reported existence of narrow vertical columns (Li et al., 2012; Ringach et al., 2016).

**Figure 5.**
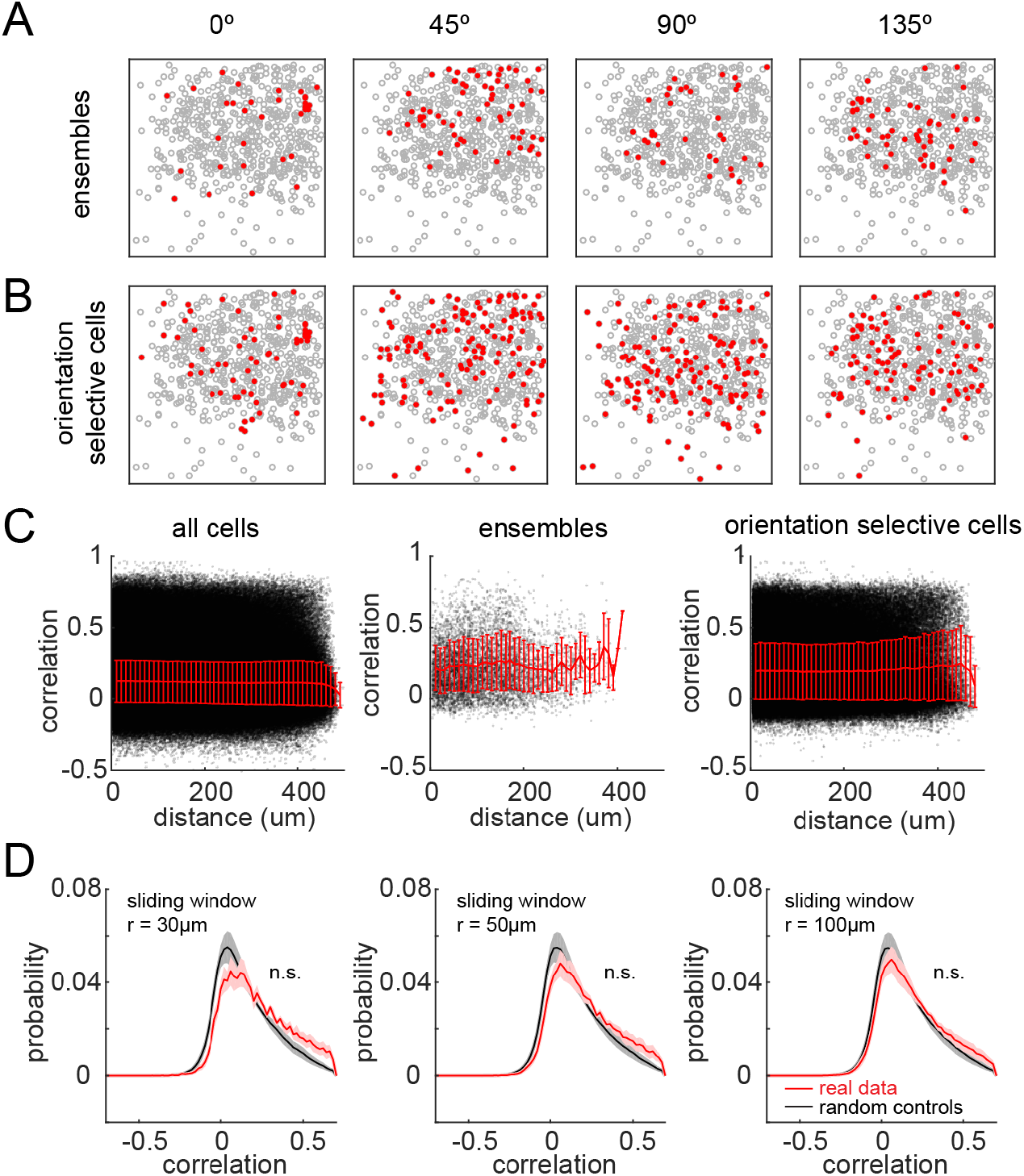
Lack of columnar structures in V1 responses. (A) Example of the spatial locations of visually-evoked ensembles (top-projection views from all planes). (B) Example of spatial locations of orientation selective cells (top-projection views from all planes). Note the salt-and-pepper structure in both cases. (C) Scatter plot of pairwise lateral distance and correlation, among all cell pairs (left), among ensembles (middle), and among orientation selective cells (right). Red line shows mean ± S.D. Data pooled from 6 experiments. (D) Distribution of pairwise correlation within columns of 30, 50 and 100 μm diameter. Red: real data; black: random controls. Random controls were generated by computing the correlation between cells within the column and a random set of cells with the same number outside of the column, repeated 50 times each column. n.s., not significant. Statistics was done by Wilcoxon rank-sum test in each correlation bin of individual datasets, comparing real data with random controls. All correlation bins above −0.3 were not significant, for all experiments. (n = 6 experiments)

### Volumetric imaging of interactions between long-range projection axons and local somas

As a final demonstration of the biological utility of our method, we sought to capture the input-output response of a circuit, by simultaneously imaging an incoming presynaptic axonal population and the responses of a postsynaptic population of cells. Indeed, the simultaneous dual-color excitation with two lasers in our system not only expands the volume that can be imaged at once through wavelength multiplexing, but also provides a tool to image and identify two distinct populations simultaneously. Using this microscope, we studied the functional interactions between the long-range axonal projections from PFC to L1 volume in V1 (labeled with GCaMP6s) and local neurons in L2/3 of V1 (labeled with jRGECO1b) (Figure 6A). We imaged the spontaneous activity of both structure with 4 planes from 25 μm to 100 μm in L1, and 4 planes from 150 μm to 300 μm in L2/3, achieving a volume rate of 13.0 vol/sec (Figure 6B). ROIs and fluorescence traces in L2/3 were extracted using CNMF algorithm as described above, and ROIs and traces in L1 were extracted using a recently developed simultaneous denoising, compression and demixing (PMD) algorithm with penalized matrix decomposition (Buchanan et al., 2018) (Figure 6C-D). The latter results in fragmented ROIs that represent putative axonal fragments and boutons with an improved SNR through denoising techniques. To group these putative ROI fragments that are potentially from the same projection, we clustered the activity traces using affinity propagation (Dueck, 2009), which does not require a cluster number input, but could identify clusters of ROIs that exhibit highly correlated activity patterns (Figure 6E, four examples shown on right). Examples of these super ROI groups are shown in Figure 6F. These ROI clusters show higher internal correlation than randomly sampled controls, indicating functional correlation (Figure 6G; n = 11 experiments; control 0.173 ± 0.005, data 0.636 ± 0.007). As the activity of long-range axonal projections and local somas are near-simultaneously recorded, we could further use the collected dataset to investigate the correlation structure of the interactions between these two populations (Figure 6H). While the overall population correlation is distributed around zero (Figure 6J; n = 11 experiments; mean correlation 0.006 ± 0.012, p = 0.617, *t*-test), we could identify ROI pairs from L1 PFC projection and L2/3 V1 soma that are highly correlated (Figure 6I), revealing functional relationships between these two populations.

**Figure 6.**
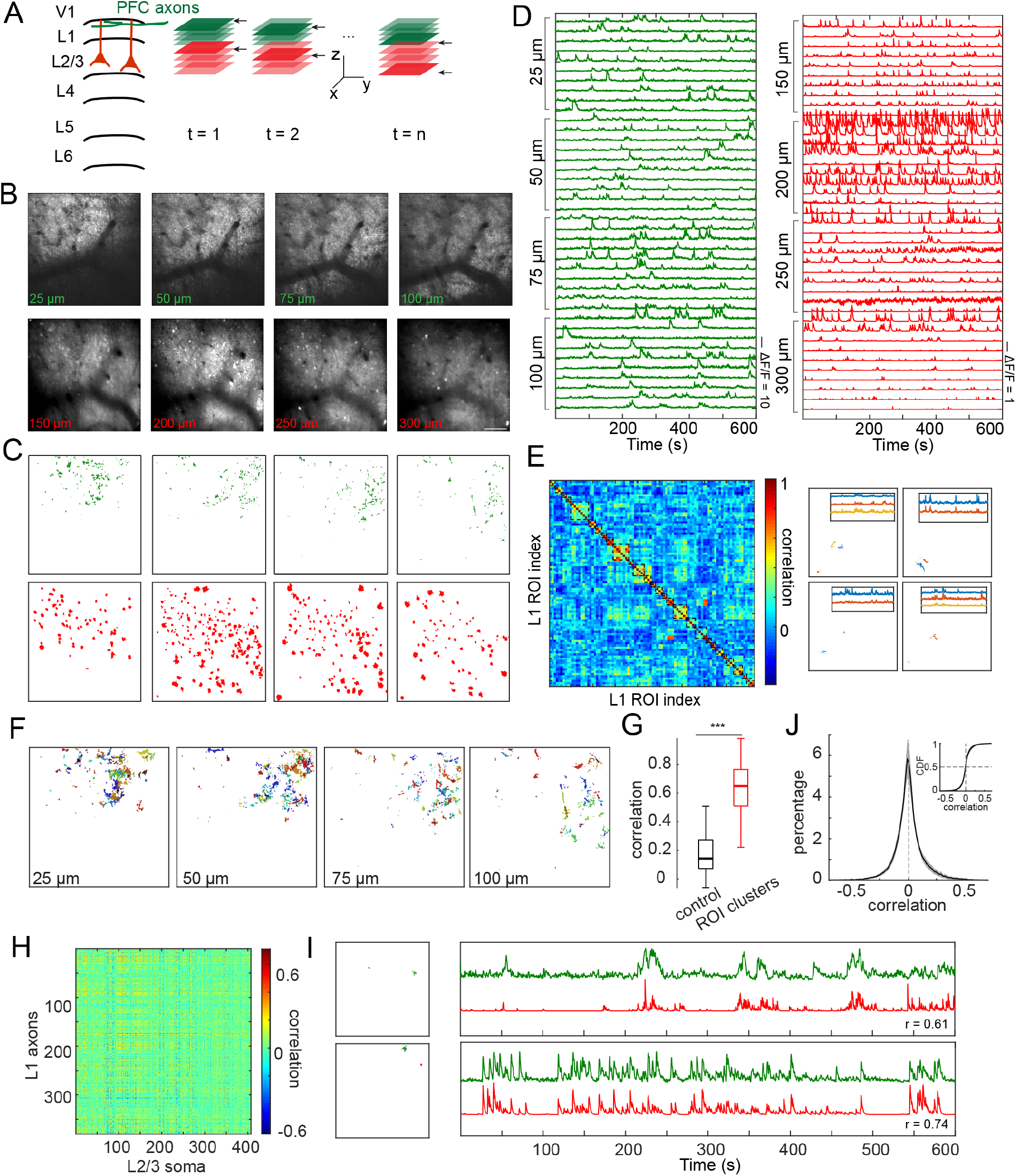
Volumetric imaging of layer 1 long-range axonal projections from PFC and local layer 2/3 somas. (A) Schematic of experiment design. In this experiment, the 920 nm laser path covers 4 planes in layer 1, and the 1064 nm path covers 4 planes in layer 2/3. (B) Examples of average images from the recorded planes, from 25 μm to 100 μm with a spacing of 25 μm in layer 1, and from 150 μm to 300 μm with a spacing of 50 μm in layer 2/3. Scale bar: 100 μm. (C) Examples of extracted ROIs in each plane. (D) Example traces from each plane. (E) Example of clustered correlation matrix of all ROIs in layer 1, sorted by cluster identity (left), and four examples of ROIs that are identified in the same clusters with corresponding traces (right). Note that the imaging depth of the example ROIs on the right might be different. (F) Clustering result in each plane. ROIs belonging to the same cluster are shown with the same color. (G) Correlation within clusters, compared with random controls (p<0.001). (H) Example of correlation between layer 1 ROIs and layer 2/3 ROIs. (J) Distribution of correlation between layer 1 and layer 2/3 ROIs. Upper right curve shows the cumulative distribution. (n = 11 experiments; mean correlation 0.006 ± 0.012, p = 0.617, *t*-test). (I) Two examples of ROI pairs from layer 1 (green) and layer 2/3 (red) that are highly correlated. The lateral locations of ROI pairs are shown on the left, and their traces shown on the right.

Similarly, our system also enables volumetric imaging of L1 apical dendrites with L2/3 somas from the same population at the same time. To do so, we labeled the V1 population with a co-injection of GCaMP6s and jRGECO1b, and simultaneously imaged the spontaneous activity of apical dendrites with the green path with 4 planes from 25 μm to 100 μm, and somas with the red path with 4 planes from 150 μm to 300 μm, at a volume rate of 13.0 vol/sec (Figure S3A-B). ROIs and traces were extracted as above (Figure S3C-D). The correlation between L1 and L2/3 is distributed slightly higher than zero (Figure S3E-F; n = 10 experiments; mean correlation 0.061 ± 0.011, p < 0.001, *t-*test), indicating a stronger spontaneous functional correlation within the same local population.

## Discussion

### Volumetric imaging through wavelength multiplexing

In this work, we extended the conventional two-dimensional two-photon imaging system to a three-dimensional volume through wavelength multiplexing, combining dual color excitation and emission with ETL/SLM-based fast z-scan approaches. By using a red calcium indicator, jRGECO1b, that reduces the loss of photons from tissue scattering, and an SLM as a z-scan device that can also implement adaptive optics to correct system aberration, we optimized our system for deep layer imaging. We demonstrated successful volumetric calcium imaging *in vivo* of 10 planes at 10.4 vol/sec, spanning across layer 2/3 to layer 5, as well as 8 planes imaging at 13.0 vol/sec of layer 1 dendritic or axonal activity with layer 2/3 somatic activity. To our knowledge, our approach currently provides the most extensive method to sample a large number of neurons per second across cortical columns (up to ∼21,000 total sample/sec over 10 planes across a depth of 450 μm, with a field of view of 500 μm × 500 μm per plane). Further improvement can be expected with optimization of virus transfection. Our approach, introducing excitation-wavelength-multiplexing to parallelize the scanning process, provides a new alternative to the volumetric imaging toolbox.

Comparing with current volumetric imaging techniques that depends on a single defocusing strategy such as ETLs (Grewe et al., 2011) and remote focusing systems (Botcherby et al., 2012), our approach is equipped with two independent modules that covers two separate volumes simultaneously, thus expanding the recorded volume to twice as much as the single strategy based systems. These two volumes are separated by their excitation and emission properties, making them truly independent of each other. Comparing with our recent SLM-based holographic multiplexing approach (Yang et al., 2016), the current approach has the same advantage of simultaneously recording from two separate planes, but also extends the recorded volume while keeping both volumes at the best performance range for each z-scan device (within ± 200 μm for both), avoiding the performance decay at higher defocus planes. Additionally, through wavelength multiplexing, depth information is encoded by wavelength, and the simultaneously-recorded dual planes are collected by two separate PMTs, avoiding the post-hoc source separation and signal demixing problem. Comparing with another multiplexing approach, temporal multiplexing, where the laser beam is split into multiple beams with their laser pulses interleaved in time and focused at different positions (Cheng et al., 2011; Stirman et al., 2016), our approach does not require complex data acquisition scheme. Our approach also makes full use of both lasers, and is therefore more effective when imaging a large number of planes. We note many of these techniques are not mutually exclusive and that the wavelength multiplexing scheme can be combined with other volumetric imaging approach to further increase the imaging throughput.

Due to the excitation spectrum overlap of GCaMP6 and RCaMP or jRGECO, it has been demonstrated that a single laser, tuned between 1020 nm and 1030 nm, can be used to simultaneously image GCaMP6 and RCaMP (Inoue et al., 2015). However, since the GCaMP excitation efficiency is best around 940 nm, and RGECO/RCaMP between ∼1060 nm and ∼1150 nm (Dana et al., 2016), using a single laser to excite both indicators would compromise the fluorophore efficiency for both indicators. Our system, through two lasers set at 920 nm and 1064 nm respectively, optimizes the fluorophore performance for simultaneous dual-color imaging.

### Simultaneous recording of large neuronal population for studying single-trial dynamics

The mammalian cortex is organized into six layers, and sensory information is transformed through the interaction between different layers (Constantinople and Bruno, 2013; Douglas and Martin, 2004). Conventionally, to study the cortical dynamics in different layers or different cortical regions during sensory perception or behavioral tasks, a standard approach is to record from each layer or region of interest during repetitions of the task trials, then align the neuronal activity with the trial start (Allen et al., 2017; Chen et al., 2017; Heindorf et al., 2018). Although trial structures provide an important reference for the underlying cortical activity, the nervous system is intrinsically noisy and variable, and it is still challenging to study the correlation structure between layers or areas with non-simultaneous recordings. One advantage of our volumetric imaging system is we could simultaneously record from a large population across multiple layers until layer 5, providing an important tool to study cross-layer computation. As demonstrated in Figure 4, simultaneous volumetric imaging reveals a distinct layer-layer correlation structure than that could be described using separated trials. We expect that our system will provide a powerful tool for studying cortical-cortical interactions in the future.

### Lack of orientation columns in mouse V1

As a proof of the utility of the method to reveal spatial interaction in functional responses, we analyzed the correlational structure of the orientation response across layers. This is a controversial issue, since the original description of unstructured orientation responses in mouse primary visual cortex (“salt and pepper” patterns of orientation) (Ohki and Reid, 2007; Ohki et al., 2005)., has been questioned by reports of the existence of clonally related neurons that are arranged in narrow vertical strips and that have similar orientation responses (Li et al., 2012; Ringach et al., 2016; Yu et al., 2009). With our volumetric method, we could, as a third party, independently examine the validity of these claims. In our analysis, however, we find no statistically significant vertical correlations in the orientation responses. While we did not ascertain the clonal relation among neurons, and we cannot comment on these data are in principle inconsistent with the presence of vertical narrow “minicolumns” of orientation and suggest that orientation selectivity is map in 3D also in a disorganized fashion, in primary visual cortex of the mouse. Our method and analysis could be extended to the study of the spatial structure of other functional properties in the cortex or other neural circuits.

### Imaging the interaction between pre and postsynaptic populations

Besides doubling the recorded volume that could be simultaneously imaged, our dual-color design goes beyond the conventional volumetric imaging, and provides a tool to image distinct neuronal population labeled with different colors, at the same time. This includes examples of layer 1 long-range projections from other regions with local somas (Figure 6), as well as excitatory neurons with interneuron populations. Combined with the volumetric imaging ability, our system provides a tool for studying the interaction of large population of distinct subnetworks and to functionally dissect out the input-output properties of neural circuits.

In closing, we present a novel volumetric imaging method that combines wavelength and spatial multiplexing of light. As the study of neural circuits becomes increasing more sophisticated, it is likely that there will not be a single “one shoe fits all” method to functionally dissect the interactions between many different types of neurons. Instead, we imagine a hybrid future, where different methods and probes and analysis could be flexibly combined, and be properly targeted to the specific question of study.

## Acknowledgements

We thank Darcy Peterka for initial discussions, Reka Letso for help with injections, and other members of the Yuste lab for help. This work is supported by the NEI (DP1EY024503, R01EY011787). This material is based upon work supported by, or in part by, the U. S. Army Research Laboratory and the U. S. Army Research Office under contract number W911NF-12-1-0594 (MURI). S. H. is a Howard Hughes Medical Institute International Student Research fellow. W. Y. holds a career award at the scientific interface by Burroughs Wellcome Fund.

## Author Contributions

Conceptualization, S.H., W.Y. and R.Y.; Methodology, S.H. and W.Y.; Investigation, S.H. and W.Y.; Formal analysis, S.H. and W.Y.; Resources, S.H., W.Y., R.Y.; Writing - original draft, S.H.; Writing - review and editing, S.H., W.Y., and R.Y.; Funding acquisition, S.H., W.Y. and R.Y.; R.Y. assembled and directed the team and secured equipment and additional funding. All data are archived at the NeuroTechnology Center at Columbia University.

## Declaration of Interests

R.Y. is listed as inventor of the following patent: ‘Devices, apparatus and method for providing photostimulation and imaging of structures’ (United States Patent 9846313). R.Y. and W.Y. are listed as inventors of the patent application: ‘Simultaneous multiplane volumetric imaging system method to image in volume simultaneously’ (2015, USPA provisional to be announced).

## Methods

### Experimental models

Experiments were performed on C57BL/6 wild-type mice, on both males and females. Experimental animals were typically postnatal (P) day P60-P120 at the time of experiments. Animals were housed on a 12h light-dark cycle with food and water ad libitum. All experimental procedures were carried out in accordance with the US National Institutes of Health and Columbia University Institutional Animal Care and Use Committee.

### Virus injection and surgery

Virus injection was performed between P30 and P60. For virus injection, a mixture of 200 nl AAV9.hSyn.GCaMP6s.WPRE.SV40 (UPenn Vector Core) and 700 nl AAV1.Syn.NES-jRGECO1b.WPRE.SV40 (UPenn Vector Core, 19279) was injected into both layer 2/3 and layer 5 on left V1 (from lambda: X = −2500, Y = 500, Z = -250/-500 μm, 400/500 nl per site). Virus was injected with glass micropipets, at a rate of 80 nl/min. For PFC injections, 700 nl AAV1.Syn.NES-jRGECO1b.WPRE.SV40 with 200 nl buffer was injected at the same location on left V1 (Z = −300 μm) between P30 and P60; two weeks after the GCaMP injection, 400 nl AAV9.hSyn.GCaMP6s.WPRE.SV40 with 200 nl buffer was injected into left PFC (from bregma: X = 300, Y = 500, Z = −900 μm).

Approximately 4-6 weeks after the initial injection, headplate implementation and craniotomy surgery were performed on the mice. Mice were anesthetized with isoflurane (1%-2%), injected with dexamethasone (2 mg/kg body weight, subcutaneous), enrofloxacin (4.47 mg/kg, subcutaneous), and carprofen (5 mg/kg, intraperitoneal). A custom made titanium headplate was mounted on the skull centered on V1 using dental cement. A 2 mm diameter circular cranial window was made around the injection site on left V1 with a dental drill, and the cranial window was covered by a 3 mm circular glass coverslip, sealed with cyanoacrylate adhesive. The mice were allowed to recover for at least one day before experiment, and were habituated with head-fixation prior to experiments. Mice were monitored and given analgesics (5mg/kg carprofen intraperitoneal) for two days post-procedure.

### Visual Stimulation

Visual stimuli were generated using MATLAB and the Psychophysics Toolbox (Mathworks) and displayed on a monitor (Dell; P1914Sf, 19-inch, 60-Hz refresh rate) positioned 28 cm in front of the right eye. Each animal was presented two consecutive visual stimulation sessions, each session with 15 trials, and each trial with a random order of 8 drifting gratings separated by 45º. In each trial, drifting gratings (100% contrast, 0.04 cycles per degree, 2 cycles per second) were shown for 4 seconds, followed by a 6-second interval with mean luminescence gray screen.

### Dual-color volumetric imaging microscope

The microscope is designed as shown in Figure 1. Two excitation lasers were used: a tunable Ti:Sapphire laser (Chameleon Ultra II, Coherent) tuned to 920 nm with a maximum output power of ∼1.6W (140-fs pulse width, 80-MHz repetition rate), and an amplified fiber laser (Fianium) with a fixed wavelength at 1064 nm with a maximum output power of ∼6W (200-fs pulse width, 80-MHz repetition rate). Each laser power is controlled with separate Pockels cells: a Conoptics EO350-160-BK Pockels cell with a 275 driver for 920 nm laser, and a Conoptics EO350-105-BK Pockels cell with a 302 RM driver for 1064 nm laser. For 920 nm path, the beam is first expanded with a 1:7.5 telescope (focal length f1 = 40 mm, f2 = 300mm). Then, the beam passes an ETL (EL-10-30-C-NIR-LD-MV, Optotune), and is rescaled by a 4:1 telescope (f3 = 400 mm, f4 = 100 mm). For the 1064 nm path, an l/2 λ waveplate (Thorlabs; AHWP05M-980) is used to rotate the laser polarization, and the beam is expanded with a 1:4 telescope (f5 = 100 mm, f6 = 400 mm) to fill the active area of SLM. Then, the focal plane is delayed with an offset lens set [composed of two lenses (f = 500 mm, −100 mm) that contact together] with an equivalent offset of ∼200 μm at imaging plane. The beam is relayed by a 1:1 telescope (f7 = 200 mm, f8 = 200mm) before being modulated by an SLM (Meadowlark Optics; HSP512-1064; 7.68 × 7.68-mm^2^ active area, 512 × 512 pixels). The beam is then rescaled by a 3:1 telescope (f9 = 300 mm, f10 = 100 mm). Then, both beams are combined through a dichroic mirror, scanned first by a resonant scanner (CRS 8K resonant scanning system), then delayed by a telescope that is composed by two equivalent lens complexes (Fig. 1B) (Stirman et al., 2016) installed in the opposite direction, then scanned by a galvanometric scanner (6215HM40B, Cambridge Technology). Both scanners are positioned at the conjugate plane to the objective pupil. The scan lens (Olympus pupil transfer lens, fscan = 50 mm) and tube lens (ftube = 180 mm) are from a modified Olympus BX-51 microscope. Imaging was done with an Olympus 253 N.A. 1.05 XLPlan N objective. The single frame rate is 60 Hz (256×256 pixels). Emission fluorescence was collected through two separate photomultiplier tubes (PMTs; Hamamatsu; H7422P-40) and two low noise amplifiers (FEMTO DHPCA-100), with a collection bandpass filter of 510 ± 40 nm (Chroma, ET520/40m) for the green path, and a 630 ± 75 nm bandpass filter (Chroma, ET630/75m) for the red path. ScanImage 2016 (Pologruto et al., 2003) was used to control the Pockels cells, the focus of the ETL and SLM, the scanning mirrors and the digitizer for data storage. Locomotion of the animals was recorded with an infared LED/photodarlington pair (Honeywell S&C HOA1877-003), which consists of a small c-shaped device positioned at the edge of the rotating wheel (striped with black tape) connected to the imaging computer as an analogue input. Locomotion was detected as voltage changes in the photodarlington readout. The typical imaging power ranges from ∼10 mW to ∼200 mW, depending on the depth.

### Adaptive optics

As SLM is a natural choice for correcting wavefront aberration for both system and samples, the excitation efficiency of the SLM path in our system can be improved by implementing adaptive optics (AO) through SLM (Ji et al., 2012; Love, 1997). Here we implemented system correction by modeling the wavefront aberration with the first 30 modes of Zernike polynomials. This includes common aberrations such as spherical aberration, astigmatism, coma, etc. We measured the coefficient for each Zernike polynomial using 0.5 μm fluorescent beads, and the final correcting wavefront on SLM is a combination of weighted Zernike polynomials with measured coefficients.

### Image processing and signal extraction

The raw imaging datasets were first motion corrected using an ImageJ plugin Moco (Dubbs et al., 2015). All imaging planes in the same datasets were registered using the same motion profile estimated from the most representative plane. Then, for somatic imaging datasets, putative neuronal regions of interest (ROIs) were initialized manually by playing through each plane of the datasets and generating a list of centroid locations using an ImageJ plugin Time Series Analyzer, in order to obtain an accurate guess of cell locations. The ROIs were then segmented by a modified version of a constrained nonnegative matrix factorization (CNMF) algorithm (Pnevmatikakis et al., 2016) that initializes with the manual list of ROI locations, and the algorithm automatically estimates the raw fluorescence signals, the denoised (filtered) signals, and the deconvolved signals. Then, all ROIs are manually selected using a custom Matlab GUI that displays both the shape of ROIs and the corresponding traces. ROIs that exhibit reasonable shape and active (not silent through the entire imaging session) were kept.

To remove potential duplicated cells that show up in adjacent planes or contaminated from the other imaging channel, ROI pairs that (1) have a Pearson correlation coefficient higher than 0.75, (2) are within 30 μm apart laterally, and (3) are within 50 μm apart axially (potential contamination from adjacent planes), or (4) come from simultaneous recorded dual planes of two channels, were kept only the ROI with highest signal-to-noise ratio (SNR).

For dendritic and axonal imaging datasets, a penalized matrix decomposition (PMD) algorithm was used to automatically denoise and demix the datasets, which improved the resulting SNR for noisy dendritic/axonal imaging (Buchanan et al., 2018). Then, dendritic or axonal ROIs were automatically segmented, and fluorescence traces were extracted by the algorithm. The traces were then filtered by trend filtering as described in the above reference. After that, ROIs were manually selected using the custom Matlab GUI as described above.

### Orientation tuning analysis

Orientation tuning curves were calculated by averaging the ΔF/F response traces of all grating stimulus sessions. This gives the polar plots. Orientation selectivity indices were calculated using circular statistics, defined as OSI = |∑_*k*_*R_k_*exp(2*iθ_k_*) /∑_*k*_*R_k_* |, where R_k_ is the response to each orientation (k = 1-8), *i* is the imaginary unit, and *θ_k_* is the orientation in radians (Tzvetanov, 2016). Neurons with OSI > 0.2 were defined as orientation selective cells. The preferred orientation was determined by the orientation that evoked the strongest ΔF/F response.

### Ensemble identification

Ensembles were detected using a graph-based community detection method, the Louvain method (Blondel et al., 2008). This method aims at detecting community structures in graphs, which are subsets of highly interconnected nodes. To apply this method, we first computed the pairwise similarity matrix using the inferred (deconvolved) fluorescence traces. Running epochs were excluded in order to reduce correlation artifact. This results in a *N*_neuron_-by-*N*_neuron_ correlation matrix. Then, to further reduce noise, weak correlation values that are below mean + 3S.D. were zeroed. A Matlab module was used to perform Louvain community detection (Rubinov and Sporns, 2010). This method does not require a cluster number input, however, a resolution parameter γ was used to control the size of output communities, with γ = 1 resulting in classic communities, γ < 1 detecting larger communities, and γ > 1 detecting smaller communities. We ranged γ between 1 and 1.5 with an interval of 0.05, performed community detection with each γ, and cross-validated using the visual stimulus prediction performance of the resulting communities, taking the γ that gives best overall prediction performance. To calculate visual stimulus prediction performance, we considered the detected communities as *N*_neuron_-by-1 population vectors, where entries corresponding to the constituent neurons in the communities are 1, and others are 0. We computed the cosine similarity between these community population vectors and real data, and used the output similarity values to compute the standard receiver operating characteristic (ROC) curves and the area under curve (AUC). AUC = 0.5 represents chance level, while larger than 0.5 represents positive predictions, and smaller than 0.5 represents negative predictions. For each detected community, we summed the AUC values for each grating direction, and combined the opposite directions to be a single orientation. For each orientation, communities that have an average AUC higher than 10% above chance level (0.55) were considered to be visually-evoked ensembles.

### Clustering of axonal activity

We used affinity propagation to cluster axonal activity in layer 1 (Dueck, 2009). This method operates on the pairwise similarity matrix between all pairs of data points, and identify the exemplers based on an input preference vector, then automatically determines the number of clusters. The preference vector was set to 95% quantile of the similarity matrix here. The Matlab module used is available at [https://www.psi.toronto.edu/index.php?q=affinity%20propagation] (without sparsity).

---

**Supplemental Figure 1.**
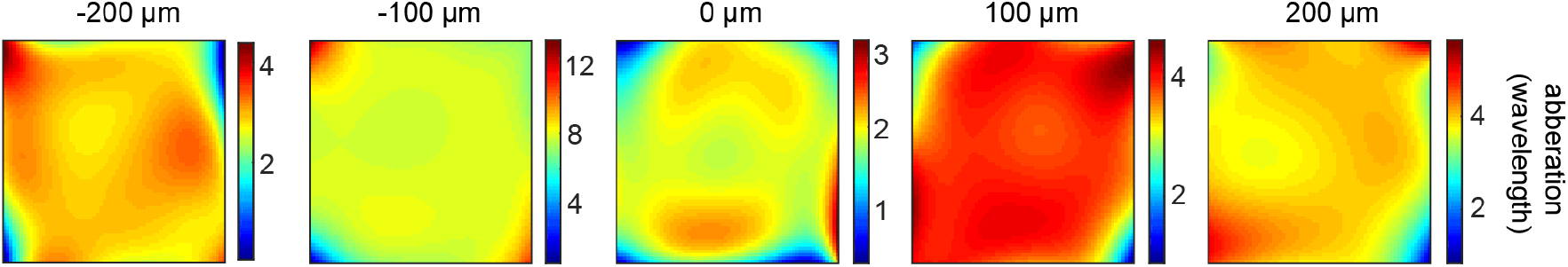
Corrective wavefront of SLM imaging path using adaptive optics. Final corrective wavefront compensated by SLM using adaptive optics, at −200 μm, −100 μm, 0 μm, 100 μm, and 200 μm SLM defocus.

**Supplemental Figure 2.**
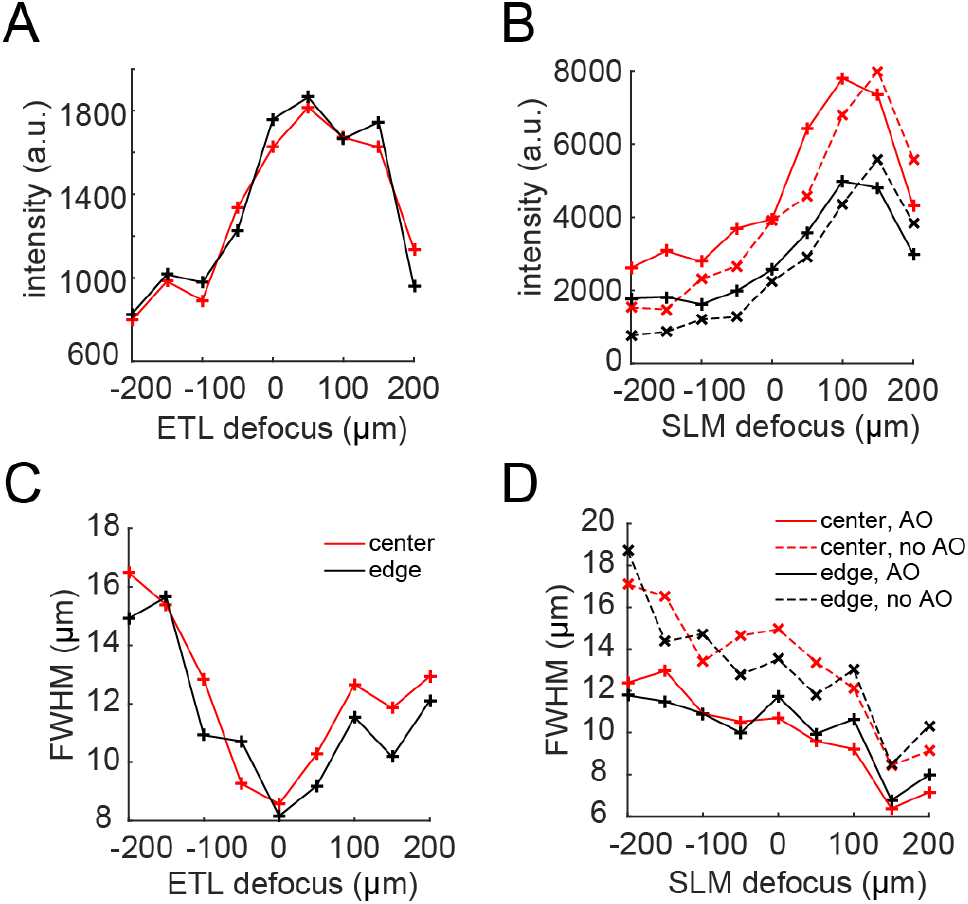
System characterization. (A) Intensity profile of the ETL path over a defocus range of −200 μm to 200 μm, measured at the center and edge of the field of view (FOV). (B) Intensity profile of the SLM path over a defocus range of −200 μm to 200 μm, measured at the center and edge of the FOV, with or without adaptive optics (AO). (C) Full-width-at-half-maximum (FWHM) of the ETL path, measured at the center and edge of FOV. (D) FWHM profile of the SLM path, measured at the center and edge of FOV, with or without AO. Note that AO improves both intensity profile and FWHM for the SLM path.

**Supplemental Figure 3.**
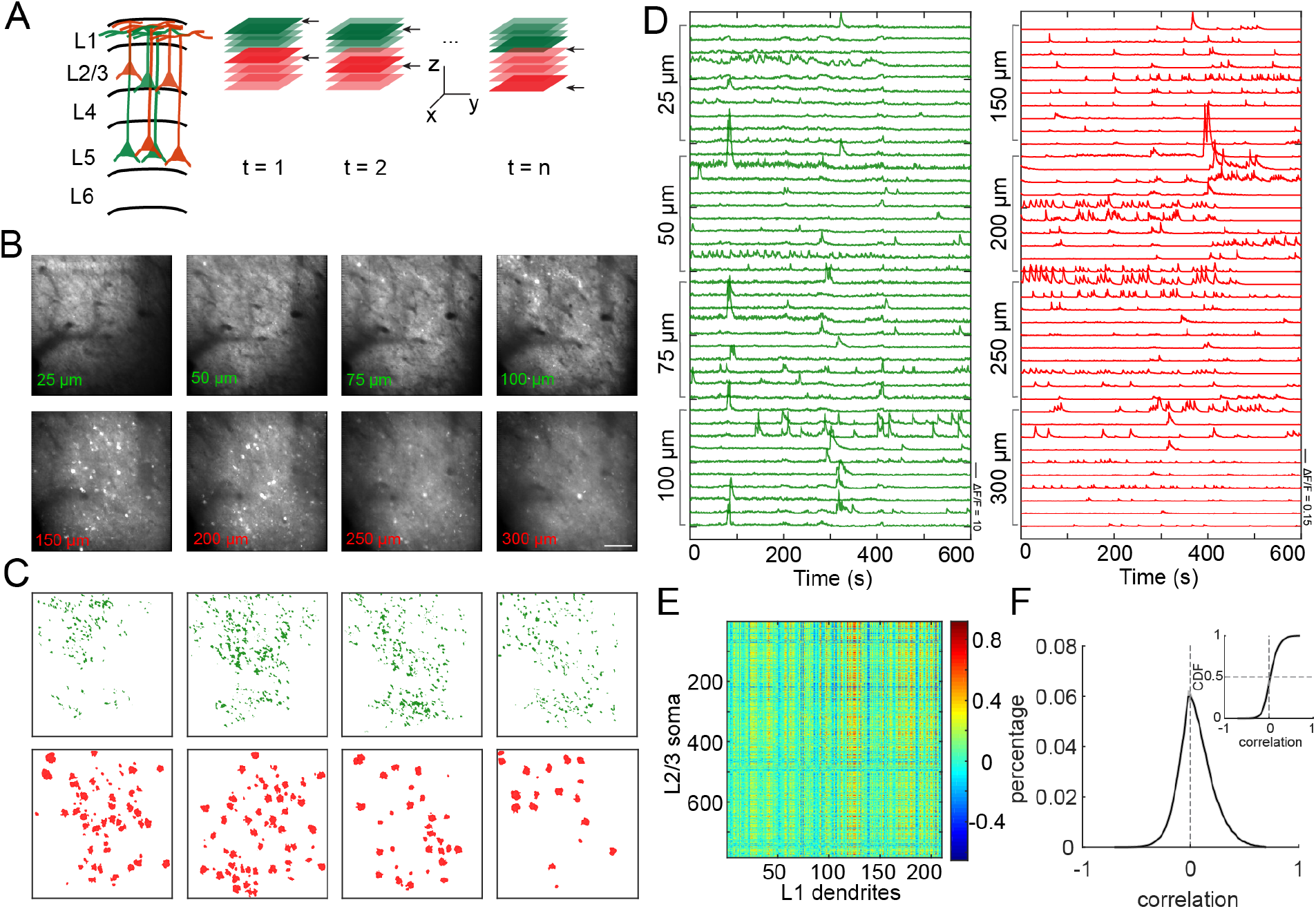
Volumetric imaging of layer 1 apical dendrites and layer 2/3 soma in V1. (A) Schematic of experiment design. The 920 nm laser path covers 4 planes in layer 1, and the 1064 nm path covers 4 planes in layer 2/3. (B) Examples of average images from the recorded planes, from 25 μm to 100 μm with a spacing of 25 μm in layer 1, and from 150 μm to 300 μm with a spacing of 50 μm in layer 2/3. Scale bar: 100 μm. (C) Examples of extracted ROIs in each plane. (D) Example traces from each plane. (E) Example of correlation between layer 1 ROIs and layer 2/3 ROIs. F) Distribution of correlation between layer 1 and layer 2/3 ROIs. Upper right curves shows the cumulative distribution. (n = 10 experiments; mean correlation 0.061 ± 0.011, p < 0.001, t-test).

